# Holistic profiling of the venom from the lethal spider *Phoneutria nigriventer* by combining high-throughput ion channel screens with venomics

**DOI:** 10.1101/2022.11.17.516848

**Authors:** FC Cardoso, AA Walker, GF King, MV Gomez

## Abstract

Spider venoms are a unique source of bioactive peptides, many of which display remarkable biological stability and neuroactivity. *Phoneutria nigriventer*, often referred to as the Brazilian wandering spider, banana spider or “armed” spider, is endemic to South America and amongst the most dangerous venomous spiders in the world. There are 4,000 envenomation accidents with *P. nigriventer* each year in Brazil, which can lead to symptoms including priapism, hypertension, blurred vision, sweating, and vomiting. In addition to its clinical relevance, *P. nigriventer* venom contains peptides that provide therapeutic effects in a range of disease models. In this study, we explored the neuroactivity and molecular diversity *P. nigriventer* venom using fractionation-guided high-throughput cellular assays coupled to proteomics and multi-pharmacology activity to broaden the knowledge about this venom and its therapeutic potential and provide a proof-of-concept for an investigative pipeline to study spider-venom derived neuroactive peptides. We coupled proteomics with ion channel assays using a neuroblastoma cell line to identify venom compounds that modulate the activity of voltage-gated sodium and calcium channels, as well as the nicotinic acetylcholine receptor. Our data revealed that *P. nigriventer* venom is highly complex compared to other neurotoxin-rich venoms and contains potent modulators of voltage-gated ion channels which were classified into four families of neuroactive peptides based on their activity and structures. In addition to the reported *P. nigriventer* neuroactive peptides, we identified at least 27 novel cysteine-rich venom peptides for which their activity and molecular target remains to be determined. Our findings provide a platform for studying the bioactivity of known and novel neuroactive components in the venom of *P. nigriventer* and other spiders and suggests that our discovery pipeline can be used to identify ion channel-targeting venom peptides with potential as pharmacological tools and to drug leads.

## Introduction

Venomous animals are a highly adapted group of organisms whose evolutionary success excelled with the emergence of venom. Spider venoms, in particular, are rich in peptide knottins specialized in modulating, often with high potency and selectivity, voltage-gated ion channels that regulate the physiology of neuronal, muscular and cardiac systems (Cardoso and Lewis, 2018;Cardoso, 2020). Although such effects can be deleterious to envenomated animals, venom components can be tailored to selectively modulate ion channels in pathways of complex diseases such as chronic pain, motor neuron disease, and epilepsy. This has been demonstrated for numerous spider venoms (Smith et al., 2015;Cardoso and Lewis, 2018;2019), including the venom of the infamous South American ctenid spider *Phoneutria nigriventer*, often referred as Brazilian wandering spider, banana spider or “armed” spider (Peigneur et al., 2018). Besides its clinical relevance due to frequent envenomation cases in Brazil, with approximately 4,000 cases per year (Isbister and Fan, 2011;Gewehr et al., 2013), *P. nigriventer* venom contains peptides that have therapeutic effects in a range of disease models including chronic pain (Pedron et al., 2021;Cavalli et al., 2022), Huntington’s disease (Joviano-Santos et al., 2022), glaucoma (da Silva et al., 2020) and erectile dysfunction (Nunes da Silva et al., 2019).

Initial studies of *P. nigriventer* venom employed fractionation via gel filtration and reversed-phase chromatography to separate the venom into five distinct groups of peptides based on their molecular weight and hydrophobicity properties; these groups were named PhTx1 to PhTx5 (Peigneur et al., 2018). Groups PhTx1–4 comprise cysteine-rich peptides that are active on voltage-gated calcium (Ca_V_), sodium (Na_V_) and potassium (K_V_) channels, while PhTx5 is comprised of short linear peptides, with a total of 34 peptides identified (Peigneur et al., 2018). Proteotranscriptomic studies of *P. nigriventer* venom revealed additional peptides with high similarity to those previously described, but very few have been characterised pharmacologically (Cardoso et al., 2003;Richardson et al., 2006). This represents an obstacle to the exploration of the therapeutic potential of *P. nigriventer* venom.

Advances in venom-peptide research have yielded high-throughput cellular screens for the discovery and pharmacological characterisation of naturally occurring molecules with activity at ion channels and receptors in physiological pathways (Cardoso et al., 2015;Cardoso et al., 2021). These methods require only a small amount of venom compared to more traditional methods and allow the identification of therapeutically relevant peptides in the early stages of the screening. Besides drug development applications, these same bioassays can assist in unravelling the bioactivity of crude and fractionated venoms from biomedically relevant venomous animals to support studies of evolution and antivenom development, but much work remains to be done in this field.

This study aimed to provide a proof-of-concept in applying high-throughput cellular screens for multiple neuronal ion channels along with proteomic studies of fractionated venom to rapidly characterise spider venoms in terms of bioactive components. It was anticipated that such a pipeline would support envenomation and evolutionary studies and the development of therapeutics from animal venoms. The venom of *P. nigriventer* (Figure 1A) was selected as a model system due to its medical relevance, the considerable number of therapeutically relevant peptides already uncovered in the venom, and the wide knowledge base available. Our approach enabled identification of potent modulators of voltage-gated ion channels which were classified into four families of neuroactive peptides based on their activity and structures. In addition to the previously characterised neuroactive peptides in the *P. nigriventer* venom, we identified 27 additional cysteine-rich venom peptides in which neuroactivities is underexplored. This work contributes to the on-going discovery and structure-function characterisation of spider-venom peptides. Moreover, our bioassay pipeline can be used to guide future research into the discovery of venom peptides that modulate the activity of ion channels, and their development as pharmacological tools and drug leads.

**Figure 1.**
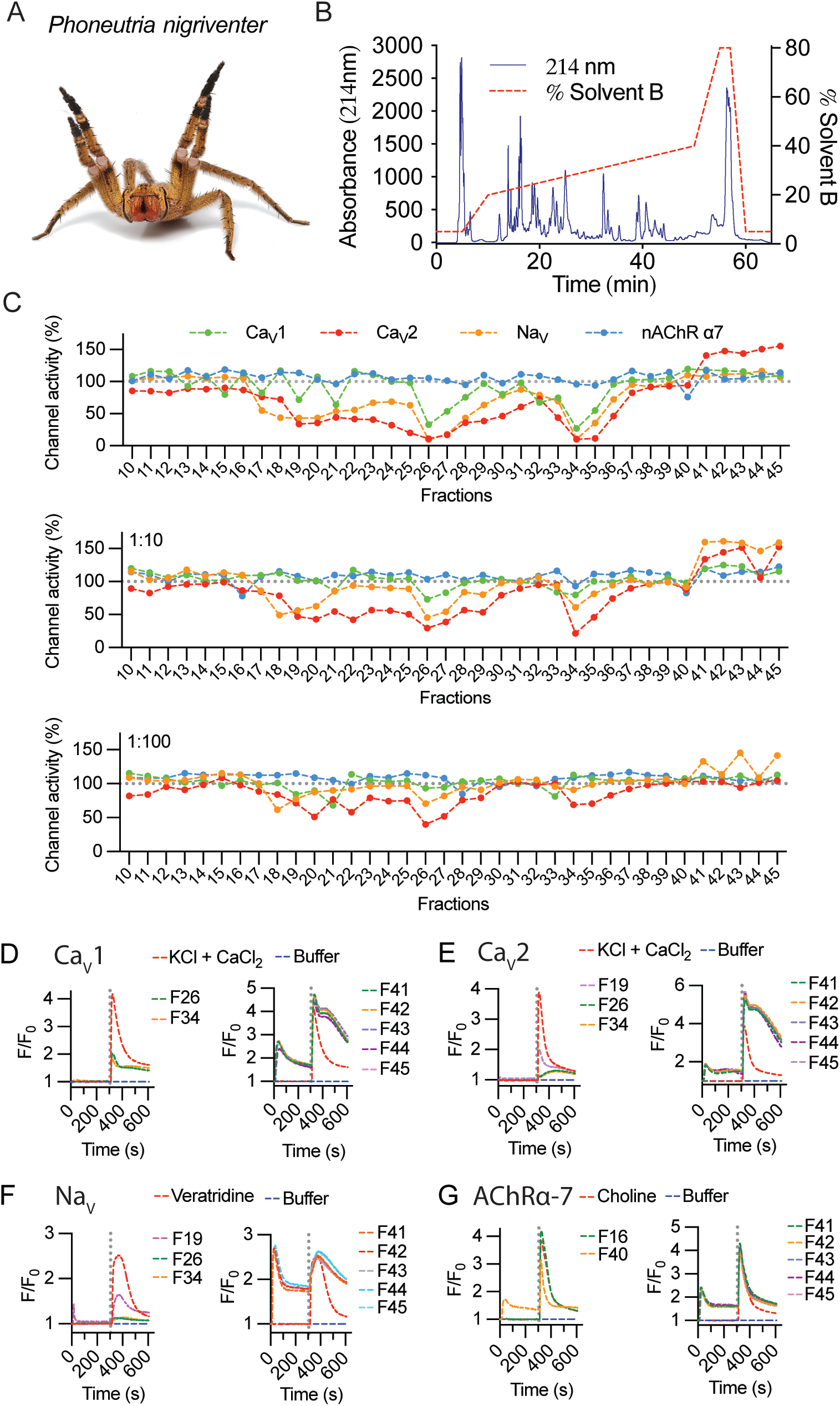
Fractionation and activity of *P. nigriventer* venom. (**A**) *P. nigriventer* specimen displaying threat posture (photo copyright Alan Henderson, www.minibeastwildlife.com.au). (**B**) RP-HPLC fractionation of 1 mg *P.nigriventer* venom. (**C**) Ion channel responses calculated from the area under the curve (AUC) after addition of selective activators for fractions 10 to 45, normalized to responses in the absence of venom fractions. (**D-E**) Representative fluorescence traces of the intracellular calcium responses of SH-SY5Y cells evoked by KCl + CaCl_2_ in the presence of venom fractions 26 and 34 for Ca_V_1, fractions 19, 26 and 34 for Ca_V_2, and fractions 41 to 45 for both Ca_V_1 and CaV2 channels. (F) Representative fluorescence traces of the intracellular calcium responses of SH-SY5Y cells evoked by veratridine and in the presence of venom fractions 19, 26 and 34 and fractions 41 to 45. (**G**) Representative fluorescence traces of the intracellular calcium responses of SH-SY5Y cells evoked by choline and in the presence of venom fractions 16 and 40 and fractions 41 to 45. Grey dotted line indicates the KCl + CaCl_2_, veratridine or choline addition.

## Methods

### Cell culture

The human neuroblastoma cell line SH-SY5Y was maintained at 37**°**C in a humidified 5% CO_2_ incubator in Roswell Park Memorial Institute (RPMI) medium supplemented with 15% foetal bovine serum (FBS) and 2 mM L-glutamine. Replicating cells were sub-cultured every 3–4 days in a 1:5 ratio using 0.25% trypsin/EDTA.

### Venom fractionation

Crude venom milked from male and female specimens of *P. nigriventer* was kindly provided by Prof. Marcus Vinicius Gomez from the Institute of Teaching and Research of Santa Casa de Belo Horizonte, Belo Horizonte, Brazil. Venom (dried, 1 mg) was dissolved in 100 μl Milli-Q water containing 0.05% trifluoroacetic acid (TFA) (Auspep, VIC, AU) and 5% acetonitrile (ACN) and centrifuged at 5,000 x g for 10 min to remove particulates. Venom was fractionated by reversed-phase high performance liquid chromatography (RP-HPLC) using a C18 column (Vydac 4.6 x 250 mm, 5 μm, Grace Discovery Sciences, USA) with a gradient of solvent B (90% ACN in 0.045% TFA) in solvent A (0.05% TFA). The gradient was 5% B for 5 min, followed by 20 to 40% solvent B over 60 min at a flow rate 0.7 mL.min^-1^. Peaks were collected every minute, with fraction 1 eluted between 1 and 2 min and so on for the other fractions. Venom fractions were lyophilised before storage at −20°C.

### Calcium influx assays

Venom fractions were screened for neuroactivity at human (h) Na_V_, hCa_V_1, hCa_V_2 and the α7 subtype of the human nicotinic acetylcholine receptor (nAChR-α7) as previously described (Cardoso et al., 2015). Briefly, SH-SY5Y cells were plated at 40,000 cells per well in 384-well flat clear-bottom black plates (Corning, NY, USA) and cultured at 37**°**C in a humidified 5% CO_2_ incubator for 48 h. Cells were loaded with 20 μl per well Calcium 4 dye (Molecular Devices) reconstituted in assay buffer containing (in mM) 140 NaCl, 11.5 glucose, 5.9 KCl, 1.4 MgCl_2_, 1.2 NaH_2_PO_4_, 5 NaHCO_3_, 1.8 CaCl_2_ and 10 HEPES pH 7.4 and incubated for 30 min at 37**°**C in a humidified 5% CO_2_ incubator. For the hCa_V_1 assay, the dye was supplemented with 1 μM ω-conotoxin-CVIF (CVIF) to inhibit Ca_V_2, and in the hCav2 assay the dye was supplemented with 10 μM nifedipine to inhibit Ca_V_1. For the hAChR-α7 assay, the dye was supplemented with PNU-120596 (Sigma-Aldrich), a positive allosteric modulator of hAChR-α7. Fluorescence responses were recorded using excitation at 470–495 nm and emission at 515–575 nm for 10 s to set the baseline, then 300 s after addition of 10% venom fraction serial diluted at 1, 1:10, and 1:100, and for a further 300 s after addition of 50 μM veratridine for hNa_V_, 90 mM KCl and 5 mM CaCl_2_ for hCa_V_, and 30 mM nicotine for hAChR-α7.

### Proteomics

Venom fractions eluting between 10 and 45 min on RP-HPLC were analysed by mass spectrometry to investigate the masses and primary structures of their peptide components. Native mass determinations were carried out with 20% of each fraction dried by vacuum centrifuge and resuspended in 20 μl 1% formic acid (FA), followed by analysis using by liquid chromatography/tandem mass spectrometry (LC-MS/MS). For identification of primary structures, 20% of each peptide fraction was reduced and alkylated by adding 40 μl of reagent composed of 4.875 mL ACN, 4.5 mL ultrapure water, 0.5 mL 1M ammonium carbonate pH 11.0, 100 μL 2-iodoethanol and 25 μL triethylphosphine, and incubating for 1 h at 37°C. Samples were speed dried in a vacuum centrifuge, and digested with 40 ng/μL trypsin in 50 mM ammonium bicarbonate pH 8.0 and 10% ACN overnight at room temperature. Trypsin was inactivated by adding 50 μL solution containing 50% acetonitrile and 5% formic acid (FA), dried in speed vacuum centrifuge, and resuspended in 1% formic acid.

LC-MS/MS samples were loaded onto a 150 × 0.1 mm Zorbax 300SB-C18 column (Agilent, Santa Clara, CA, USA) on a Shimadzu Nano LC system with the outflow coupled to a SCIEX 5600 Triple TOF (Framingham, MA, USA) mass spectrometer equipped with a Turbo V ion source. Peptides were eluted using a 30 min gradient of 1–40% solvent B (90% ACN/0.1% FA) in solvent A (0.1% FA) at a flow rate of 0.2 mL/min. For MS1 scans, *m/z* was set between 350 and 2200. Precursor ions with *m/z* 350–1500, charge of +2 to +5, and signals with >100 counts/s (excluding isotopes within 2 Da) were selected for fragmentation, and MS2 scans were collected over a range of 80–1500 *m/z*. Scans were obtained with an accumulation time of 250 ms and a cycle of 4 s.

A database of possible peptide sequences produced in *P. nigriventer* venom glands was compiled using a published venom-gland transcriptome (Diniz et al., 2018), from which open reading frames (ORFs) longer than 30 amino acids were identified and translated by TransDecoder. A list of 200 common MS contaminants was added to the translated ORFs, which was used as a sequence database to compare to mass spectral data using the Paragon algorithm in Protein Pilot 2.2 software (AB SCIEX). We report only peptides for which more than two tryptic fragments were detected with >95% confidence, or where one tryptic fragment was detected, and a secretion signal peptide was predicted by SignalP5.0.

### Molecular modelling

Venom peptides identified in this study were selected based on their structure and bioactivity, and their three-dimensional (3D) structure predicted using the AlphaFold 2 algorithm (Jumper et al., 2021). All 3D structures displayed were from unrelaxed models ranked 1 for each peptide prediction. 3D structures were visualised and analysed using PyMol (PyMOL).

### Data analysis

Fluorescence traces were evaluated using the Maximum-Minimum or Area Under the Curve values generated after addition of ion channel activator. Data were normalised against the negative control (PSS buffer control) and positive control (ion channel activator) for each assay and corrected using the response over baseline from 1 to 5 s.

## Results

### Screening of P. nigriventer venom fractions

Fractionation of 1 mg of *P. nigriventer* crude venom using RP-HPLC produced numerous peaks eluting between 20% and 40% solvent B, and fractions eluting between 11 and 45 min were selected for pharmacological analysis (Figure 1B). Screening using the SH-SY5Y neuroblastoma cell line revealed strong modulation of voltage-gated ion channels including both inhibition or enhancement of ion channel activity (Figure 1C). Venom fractions eluting between 18 and 34 min showed strong inhibition of Ca_V_ and Na_V_ activity, while fractions eluting between 41 and 45 min strongly activated Ca_v_2 channels (Figure 1C, top panel). At a dilution of 1:10, these inhibitory effects persisted for both Na_V_ and Ca_V_2 channels for fractions eluting at 19–20 min and 26–34 min, and was absent for Ca_V_1 channels (Figure 1C, middle panel). Fractions eluting from 21–25 min showed a clear preference for inhibiting only Ca_V_2 channels (Figure 1C). Interestingly, at 10:1 dilution, channel activity enhancement was stronger on Na_V_ channels compared to Ca_V_2 channels, suggesting potential concentration-dependent synergistic effects of venom peptides modulating both Na_V_ and Ca_V_2 channels. At the highest venom dilution of 1:100, persistent inhibition of Na_V_ channel was observed for fraction 20 (F20), while the remaining inhibitory fractions preferentially inhibited only Ca_V_2 channels (Figure 1C, bottom panel). Channel enhancement persisted for Na_V_ channels in fractions eluting from 41–45 min. No activity was observed against α7-nAChR at any venom concentration tested. Overall, inhibitory activity was primarily observed for fractions eluting at shorter retention times (i.e., more hydrophilic compounds), while strong ion channel activation was induced by more hydrophobic peptides with longer RP-HPLC retention times.

Fluorescent traces measured upon addition of venom fractions revealed an increase in intracellular calcium ([Ca^2+^]_i_), suggesting that these venom peptides can activate closed channels as well as enhance the responses of these channels opened using pharmacological intervention (Figure 1D-G). This was observed for Ca_V_ responses in the presence of 1 μM CVIF (Ca_V_2 inhibitor, Figure 1D) and 10 μM nifedipine (Ca_V_1 inhibitor, Figure 1E). In the absence of Ca_V_ inhibitors, these [Ca^2+^]_i_ responses resemble the levels of Ca_V_1 responses in Figure 1D as observed for F40–F45 applied in the Na_V_ channels assay (Figure 2F). The activities of inhibitory fractions were mostly free from initial [Ca^2+^]_i_ responses upon venom addition, except F19 for Na_V_ and F40 for α7-nAChR (Figure 1F and G).

**Figure 2.**
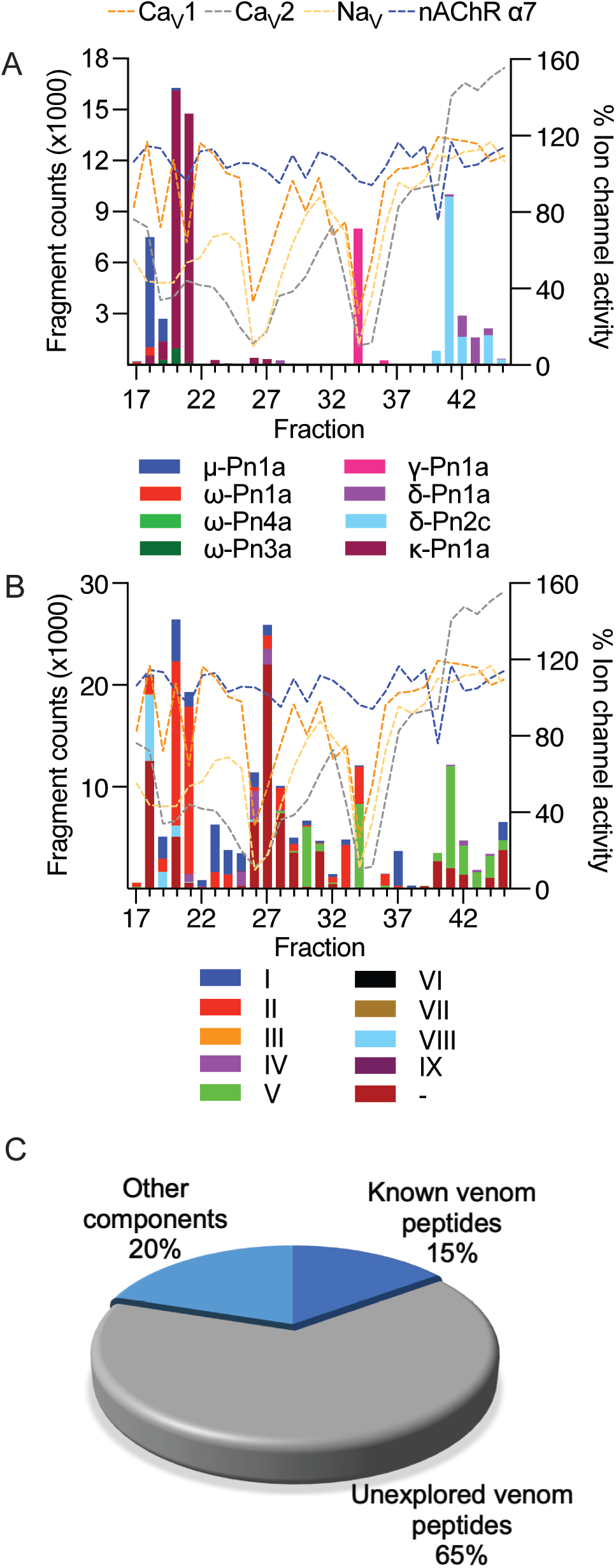
Estimated levels of peptide/protein venom components identified in fractions F17 to F45, and their respective bioactivity at Na_V_ and Ca_V_ channels and the α7-nAChR. (**A**) Venom peptides with previously reported bioactivity detected in fractions by mass spectrometry and compared to fraction bioactivity at Na_V_ and Ca_V_ channels and the α7-AChR. (**B**) Venom peptides detected in fractions classified according to their cysteine framework I to IX (Diniz et al., 2018), and compared to fraction bioactivity at Na_V_ and Ca_V_ channels and the α7-AChR. (**C**) Proportion of known and unknown venom peptides and other venom components detected in this study.

### Identification of peptides in P. nigriventer venom fractions

The venom of *P. nigriventer* has been extensively characterised in terms of composition and bioactivity (Diniz et al., 2018;Peigneur et al., 2018), including neuronal ion channel activity and proteomics, but not by using a combined approach. In this study, by combining these approaches, we were able to rapidly identify 58 peptides and proteins in the venom. Due to the complexity of previous nomenclature for *P. nigriventer* venom peptides, we refer to them here using both the rational nomenclature developed for spider toxins (King et al., 2008) and an identifying number (e.g., PN367) that is linked to a sequence and a list of previously used names in Supplementary Table S1. Of the 58 identified amino acid sequences, only eight (15%) are peptides that have had their bioactivity reported in previous studies (Figure 2A and C, Supplementary Table S1) (Peigneur et al., 2018). These included the known neuroactive components μ-CNTX-Pn1a (Tx1) (Diniz et al., 2006;Martin-Moutot et al., 2006), κ-CNTX-Pn1a (Tx3-1, PhK_V_) (Kushmerick et al., 1999;Almeida et al., 2011), ω-CNTX-Pn1a (Tx3-2) (Cordeiro Mdo et al., 1993), γ-CNTX-Pn1a (Tx4(5-5)) (Paiva et al., 2016), δ-CNTX-Pn1a (Tx4(6-1)) (de Lima et al., 2002;Emerich et al., 2016), δ-CNTX-Pn2c (Tx2-5a) (Yonamine et al., 2004), ω-CNTX-Pn4a (Tx3-6) (Cardoso et al., 2003;Vieira et al., 2005) and ω-CNTX Pn3a (Tx3-4) (Dos Santos et al., 2002) (Figure 2A). Even among these eight peptides, only a few venom peptides have had their molecular pharmacology characterized in detail (Peigneur et al., 2018), or their activities confirmed using recombinant peptides (Diniz et al., 2006;Paiva et al., 2016;Garcia Mendes et al., 2021).

Most of the identified sequences in this study (74%) represent peptides with unexplored bioactivity; 38 (65%) of the 43 peptides identified have cysteine-rich scaffolds typical of spider-venom peptides (Figure 2C). Some of these venom peptides, such as PN367 and PN363, have a type I scaffold (Diniz et al., 2018) and are predicted by Alphafold 2 to fold into cystine-knot scaffolds typical of spider-venom peptides (King and Hardy, 2013) (Figure 3). Scaffolds II-VIII either form elaborated cystine-knot folds with extra disulphide bonds, or alternative structures such as for scaffolds III and IV (Figure 3). Novel peptides with high identity with other toxins and not previously described in *P. nigriventer* venom included: PN367 displaying identity with a *Agelena orientalis* venom peptide; PN369 displaying identity with a *Lycosa singoriensis* venom peptide, and PN365 displaying scaffold III and identity with another *Lycosa singoriensis* venom peptide (Supplementary Table S1).

**Figure 3.**
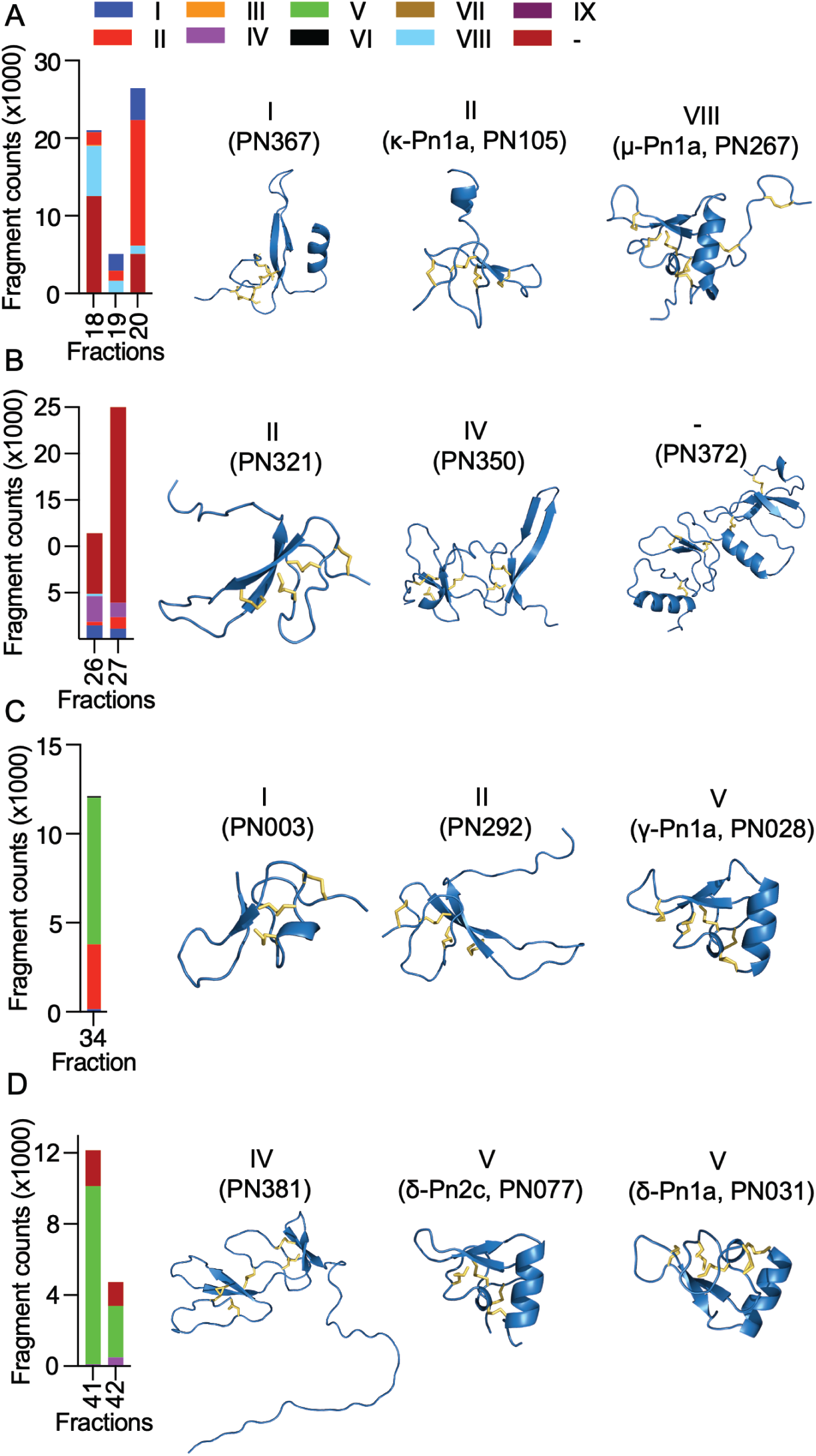
Diversity and estimated levels of cysteine-rich scaffolds identified in highly neuroactive RP-HPLC fractions from the venom of *P. nigriventer*, and their predicted 3D structures. (**A**) Fractions 18 to 20 comprised high levels of scaffolds I, II and VIII represented by the 3D structures of PN367, PN105 and PN267, respectively. (**B**) Fractions 26 and 27 comprised high levels of scaffolds II, and IV, and an undefined scaffold represented by the 3D structures of PN321, PN350 and PN372, respectively. (**C**) Fraction 34 comprised high levels of scaffolds I, II, and V represented by the 3D structures of PN003, PN292 and PN028, respectively. (**D**) Fractions 41 and 42 comprised high levels of scaffolds IV and V represented by the 3D structures of PN381, and PN007 and PN031, respectively.

Additional disulphide-rich scaffolds present in *P. nigriventer* venom include three peptides predicted by the algorithm HMMER to form a thyroglobulin type 1 repeat domain (E < e^-17^ in each case), one of which has been previously reported as U24-CNTX-Pn1a; peptide PN370 which displays high identity with a peptide found in venom of the scorpion *Scorpiops jendeki* and is predicted by the algorithm HMMER to form into a trypsin-inhibitor-like cysteine-rich domain (E < 2e^-13^); and the peptide PN376 that is predicted by HMMER to form a fungal protease inhibitor domain (E < e^-6^) (Supplementary Table S1). Additional new scaffolds identified in this study were named following the previous suggested nomenclature (Diniz et al., 2018) as X (CXCC motif, 12 Cys residues: −C−C−C−C−CXCC−C−C−C−C−C−), XI (12 Cys residues: −C−C−C−CXC−CXC−C−CXC−C−C−), XII (11 Cys residues: −C–C–CXC–CXC–C–C–CXC–C) and XIII 10 Cys residues: −C−C−C−C−C−C−CXC−C−C−), and include the peptides PN376, PN372, PN373 and PN375, and PN370, respectively.

Only 9% of the identified sequences were peptides with two or fewer Cys residues (Supplementary Table S1). F17 contained a peptide (PN361) matching a C-terminally amidated peptide precursor from *Araneus ventricosus* identified in a genomic study (Kono et al., 2019). This precursor has 70% sequence identify with the prohormone-1 like precursor from the honeybee *Apis mellifera* (UniProt P85798) which is believed to be cleaved to form three short peptides with neuronal activity. Another short peptide, PN366 identified in F18 and F28–F30, matches a neuropeptide in the sea slug *Aplysia californica* (UniProt P06518). Larger proteins were also detected in some fractions; for example, F18 and F31 contained a fragment at 58% and 70% total fraction components, respectively, matching a zinc metalloprotease from the nematode *Caenorhabditis elegans* (UniProt 55112) which contains a peptidase family M12A domain.

### Diversity of neuroactive peptides in P. nigriventer venom

The cysteine-rich scaffolds of venom peptides identified in this study were compared to the classification previously proposed for *P. nigriventer* venom peptides (Diniz et al., 2018) (Figure 2B and Figure 3). Peptides in fractions displaying inhibitory properties corresponded to scaffolds I, II, IV, V and VIII, as well as unnamed scaffolds, while peptides in fractions with activation properties comprised mostly the scaffold V. All of these scaffolds are inhibitor cystine knot motifs, except for scaffold IV which had the highest level in F26 represented by the peptide PN350.

Neuroactive peptides with greater hydrophilicity (i.e., those with short RP-HPLC retention times) showed pharmacological properties reminiscent of known spider-derived μ-toxins (F17 and F18) and ω-toxins (i.e., inhibition of Ca_V_1 and Ca_V_2 channels by F19 and F20) (Figures 1C and 4A). Major components driving those bioactivities were the pharmacologically characterised peptides μ-CNTX-Pn1a, ω-CNTX-Pn1a and ω-CNTX-Pn3a, as well as additional peptides with unknown activity (Figure 4A). As the hydrophilicity of the peptides decrease (i.e., peptides with long RP-HPLC retention times), persistent Ca_V_2 inhibition was observed with maximum inhibitory activity in F26 and F27, and with the additional peptide ω-CNTX-Pn4a detected in F24 (Figures 1C, 2A and 4B). Interestingly, venom peptides characterized as K_V_ modulators, such as κ-CNTX-Pn1a, were detected in fractions displaying strong μ- and ω-pharmacology (fractions 26 and 27); it was not clear if the observed bioactivity was also associated to the modulation of K_V_ channels.

**Figure 4.**
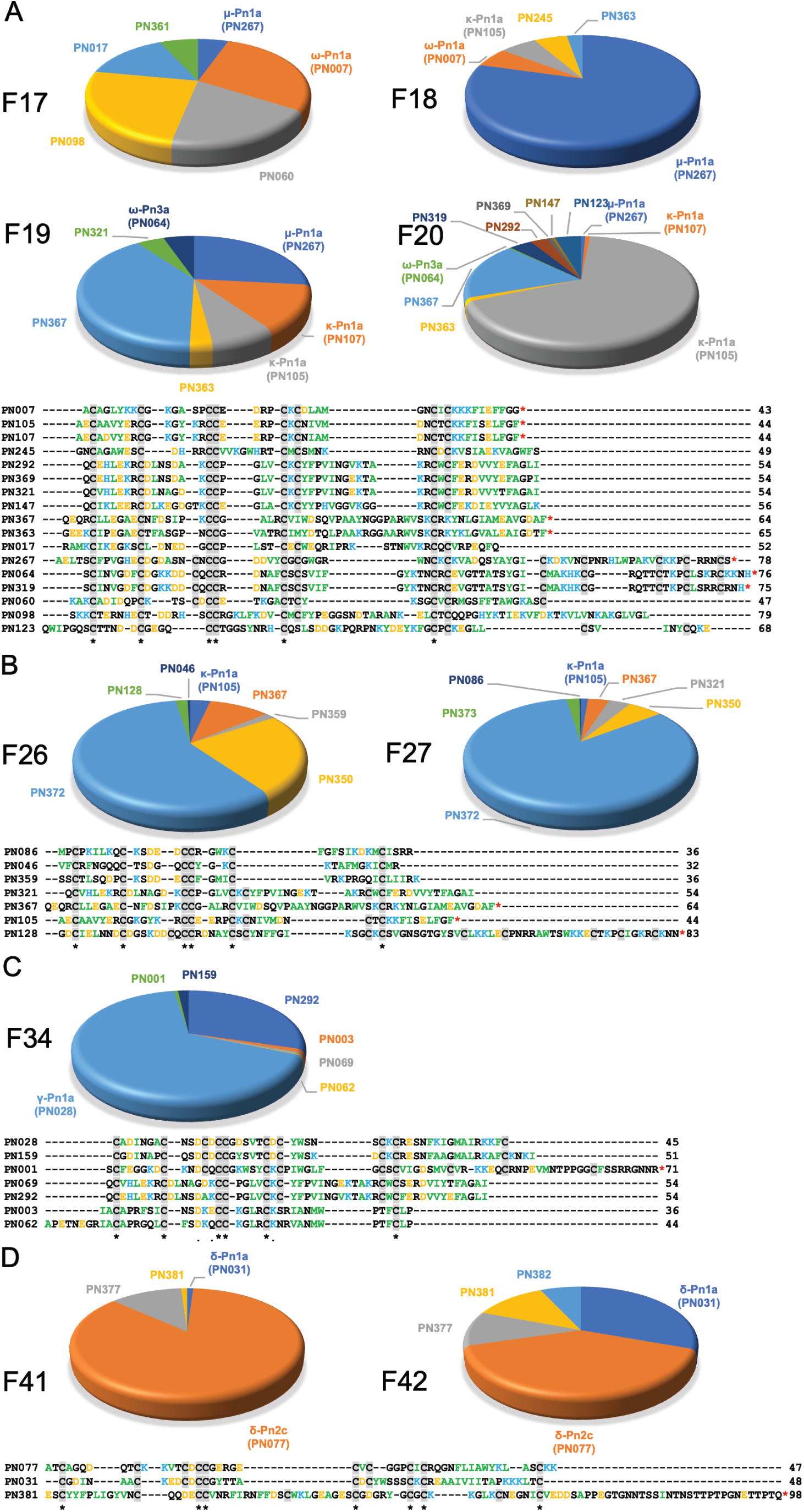
Venom peptide content of highly neuroactive RP-HPLC fractions from the venom of *P. nigriventer*. (**A**) Identification of the cysteine-rich peptides and proteins in fractions 17–20 displaying potent inhibition of neuronal Na_V_ and Ca_V_2 channels. Positively and negatively charged residues are coloured blue and orange, respectively, hydrophobic residues are green, and cysteines are highlighted in grey box. (**B**) Identification of the peptide and protein content of the fractions 16 and 27 displaying potent inhibition of neuronal Na_V_, Ca_V_1 and Ca_V_2 channels. (**C**) Identification of the peptide and protein content of the fraction 34 displaying potent inhibition of neuronal Na_V_, Ca_V_1 and Ca_V_2 channels. (**D**) Identification of the peptide and protein content of the fraction 34 displaying potent activation of neuronal NaV and CaV2 channels. Sequences labelled with a red asterisk (*) at the C-terminal are likely C-terminally amidated.

Neuroactive peptides presenting more hydrophobic structures showed properties of μ and ω-peptides, but with preference for Ca_V_2 channels as observed for fraction 34 in which the peptide γ-Pn1a is the major component, consistent with its previously observed modulation of multiple cation channels (Paiva et al., 2016); and of δ-peptides as observed in fractions 41 to 45, in which major components included the peptides δ-Pn1a and δ-Pn2c (Figures 1C-F, and Figures 4C and D). Notably, the main components of some of the most neuroactive fractions are peptides with still unexplored bioactivity, e.g. fraction 26 (Figure 2C, and Figures 3 and 4).

### Pharmacological groups and pharmacophores

Our approach allowed classification of *P. nigriventer* venom peptides into four major groups based on their bioactivity (Figure 5). **Group 1** is comprised of μ and ω peptides with scaffold type VIII and more hydrophilic properties as they eluted between F17 and F21. As representatives from this group, μ-CNTX-Pn1a and ω-CNTX-Pn3a have a potential “KR electrostatic trap”, a pharmacophore described in spider-venom peptides that modulate ion channels (Hu et al., 2021;Wisedchaisri et al., 2021), in their primary and tertiary structures (Figure 5A). This pharmacophore is likely composed of residues R61, K67, K70, K71, R74 and R75 in μ-CNTX-Pn1a and residues K54, K56, R59, K65, K70, R71, K73 and K74 in ω-CNTX-Pn3a. Within this group, the ω-CNTX-Pn3a homologue PN319 differs at three positions, making it an interesting candidate for further characterisation.

**Figure 5.**
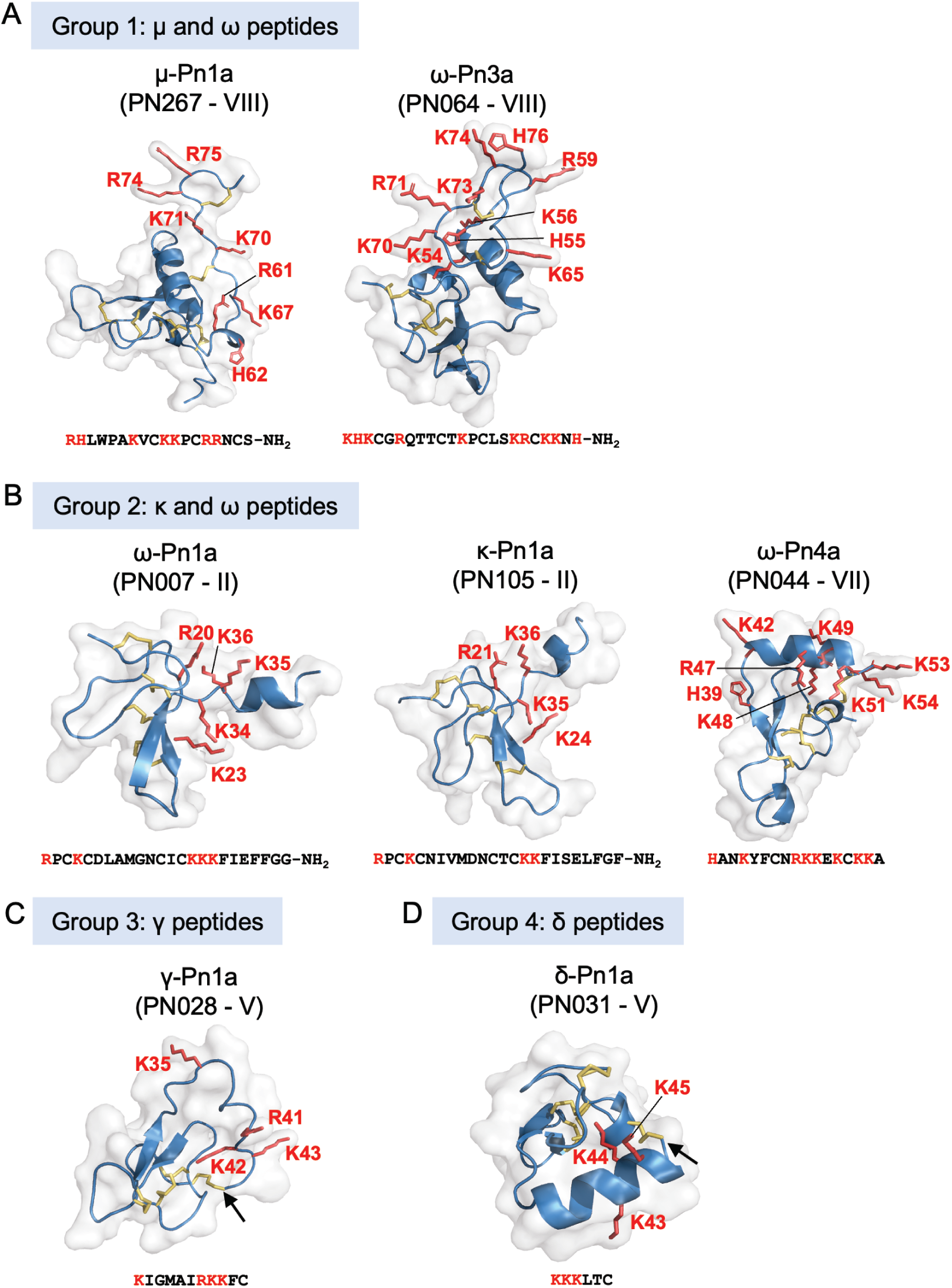
Pharmacological groups identified in the most active venom fractions highlighting the “KR electrostatic trap” pharmacophore common to spider toxins that modulate the activity of ion channels. (**A**) Group 1 is represented by μ- and ω-spider-venom peptides with large and complex type VIII scaffold. (**B**) Group 2 is represented by κ- and ω-spider-venom peptides with type II and VII scaffolds. (**C**) Group 3 is represented by γ-spider-venom peptides with type V scaffold. (**D**) Group 4 is represented by δ-spider-venom peptides displaying a type V scaffold. K and R residues located in the C-terminal region of these peptides and grouped on a positively charged face are highlighted in red in the sequences and in red tubes in the corresponding 3D structures. Arrows shows the cysteine-bridge connection forming the cyclic peptide structures predicted for PN028 and PN031.

**Group 2** comprises κ and ω peptides that eluted between F17 and F28, with scaffold types II and VII (Figure 5B). As representatives from this group, peptides κ-CNTX-Pn1a, ω-CNTX-Pn1a and ω-CNTX-Pn4a also contain a “KR trap” pharmacophore comprised of residues R20, K23, K34, K35 and K36 for ω-Pn1a; R21, K24, K35 and K36 for κ-Pn1a; and K42, R47, K48, K49, K51, K53 and K54 for ω-Pn4a. In this group, PN107 differs from κ-CNTX-Pn1a differs by only two residues and is an interesting peptide for further exploration.

**Group 3** is comprised of more hydrophobic γ peptides that eluted in F33–F36 and possess a type V scaffold (Figures 1B and 5C). It is represented by γ-CNTX-Pn1a with a potential “KR trap” comprising residues K35, R41, K42 and K43. Although γ peptides supposedly modulate non-specific cation channels, γ-CNTX-Pn1a has been reported as a β-peptide that inhibits Na_V_ channels (Paiva et al., 2016), which agrees with the results from our high-throughput ion channels assays (Figures 1C–F and 2). Interestingly, γ-CNTX-Pn1a predicted 3D structure formed a cyclic structure in which the N-terminal cysteine formed a disulphide bridge with C-terminal cysteine (Figures 3C and 5C). These same fractions contain other ICK peptides including PN003 and PN292 with scaffold types I and II, respectively; their pharmacological targets have not been explored but they likely contribute to the strong inhibition of Ca_V_ channels by F34 (Figures 1–3).

**Group 4** is composed of very hydrophobic δ peptides that elute in F40–F45 and possess a type V scaffold (Figures1B and 5D). It is represented by δ-CNTX-Pn1a with potential “KR trap” comprising residues K43, K44, and K45 (Figure 5D). In this group we also identified δ-CNTX-Pn2c which differs not only in primary structure but also in the scaffold V tertiary structure by presenting a non-cyclic structure compared to the cyclic structure predicted for δ-CNTX-Pn1a connected by the N- and C-terminal cysteines (Figures 3D and 5D). Beyond these known peptides, this group comprised interesting unexplored peptides such as PN032 and PN023 showing δ peptide domains and differing from γ-CNTX-Pn1a by 12 and 11 residues, respectively.

## Discussion

Spiders are one of the most speciose venomous taxa, with >50,000 characterised species (see https://wsc.nmbe.ch/statistics/). Their venoms are rich in neuroactive peptides that target a wide range of neuronal ion channels and receptors using mechanisms distinct from those of neurotoxins from other venomous animals such as cone snails and scorpions. The exploration of venom peptides targeting ion channels and receptors provides novel opportunities for the development of pharmacological tools to understand disease mechanisms (Cardoso and Lewis, 2018;Cardoso, 2020) as well as provision of leads for development of therapeutics (King, 2011) and bioinsecticides (Smith et al., 2013).

Spiders are classified in two major groups, or infraorders (King, 2004): Mygalomorphae, or so-called “primitive spiders”, includes the family Theraphosidae, or tarantulas, which are the most well studied spider venoms due to the large-size and long lifespan (often >20 years) of these spiders. Araneomorphae, or “modern spiders”, comprise >90% of all extant spider species, including the family Ctenidae in which *P. nigriventer* resides. Notably, despite their much greater species diversity, araneomorph venoms are underexplored compared to mygalomorphs due to their smaller size and shorter lifespan (typically 1–2 years). Our data and those of others (Binford et al., 2009;Zhang et al., 2010;Diniz et al., 2018;Peigneur et al., 2018) show a great diversity of both pharmacological actions and cysteine scaffolds in araneomorph, which may have facilitated the highly successful araneomorph radiation. This also suggest Araneomorphae’s venoms may be a rich source of unique venom peptides with more diverse structures and pharmacological functions and additional biotechnological and therapeutic applications to Mygalomorphae’s venoms.

Investigative pipelines in venomic studies often focus on the elucidation of venom components based on their structures but lack clear strategies to investigate venom bioactivities (von Reumont et al., 2022). Investigations using fractionated venom (Cardoso et al., 2015;Cardoso et al., 2017;Estrada-Gomez et al., 2019;Cardoso et al., 2021) provides more defined biological functions that is possible using crude venom due to the immense pharmacological diversity of venom, which often contains venom components with opposing activity as well as components that act synergistically s (Raposo et al., 2016). Considering the large number of extant spiders and consequently the exceptionally large number of venom components available for investigation, high-throughput (HT) functional bioassays are essential for developing a holistic understanding of venom pharmacology, and they provide a complement to venomic studies. A recent study using HT bioassays to investigate the ion channel targets of Australian funnel-web spider venoms recaptured current taxonomy and revealed potential drug targets to treat severely envenomated patients (Cardoso et al., 2022). In this present study, we also demonstrated the feasibility of applying HT functional bioassays to investigate spider venom components that mediated the activity of voltage-gated ion channels. We were able to capture all known venom components and associated bioactivities using a HT functional assay as well as several new unexplored venom peptides that warrant further exploration. This was achievable only by combining HT bioassays with transcriptomic and proteomic approaches.

The complexity of the cysteine-rich scaffolds in *P. nigriventer* venom peptides suggests that further exploration utilising recombinant or synthetic peptides might be challenging but essential, and these could also benefit from modern strategies utilizing HT recombinant expression or chemical synthesis (Pipkorn et al., 2002;Turchetto et al., 2017). In tandem with automated whole-cell patch-clamp electrophysiological studies, this will build a pipeline to further investigate known and new peptides in the venom of *P. nigriventer* and allow selection of candidates with biotechnological potential. The putative “KR trap” pharmacophores identified in those venom peptides warrants further exploration of the structure-function relationships of the diverse pharmacological groups found in the venom of *P. nigriventer*.

In conclusion, we demonstrated that the introduction of HT functional bioassays in venomic studies is essential to provide a more complete understanding of venom components in terms of structure and function. It also allows venom peptides to be ranked for further investigation based on their bioactivity and structural diversity, which is not possible via transcriptomic and proteomic studies alone. Furthermore, this study provides a guide to assist the exploration of neuroactive venoms from other animals, in particularly for the underexplored araneomorph spiders.

## Authors contribution

Conceptualization: FCC; Design, conduct, and analysis of experiments: FCC and AAW; MVG contributed with the *P. nigriventer* crude venom. Drafting of manuscript: FCC. All authors contributed to reviewing and editing of the manuscript and approved the final version for submission.

## Acknowledgements

We thank Mr Alun Jones and Dr Kuok Yap (Institute for Molecular Bioscience, The University of Queensland) for assistance with mass spectrometry experiments.

## References

Almeida, A.P., Andrade, A.B., Ferreira, A.J., Pires, A.C., Damasceno, D.D., Alves, M.N., Gomes, E.R., Kushmerick, C., Lima, R.F., Prado, M.A., Prado, V.F., Richardson, M., Cordeiro, M.N., Guatimosim, S., and Gomez, M. (2011). Antiarrhythmogenic effects of a neurotoxin from the spider Phoneutria nigriventer. Toxicon 57, 217–224.

Binford, G.J., Bodner, M.R., Cordes, M.H., Baldwin, K.L., Rynerson, M.R., Burns, S.N., and Zobel-Thropp, P.A. (2009). Molecular evolution, functional variation, and proposed nomenclature of the gene family that includes sphingomyelinase D in sicariid spider venoms. Mol Biol Evol 26, 547–566.

Cardoso, F.C. (2020). Multi-targeting sodium and calcium channels using venom peptides for the treatment of complex ion channels-related diseases. Biochem Pharmacol 181, 114107.

Cardoso, F.C., Castro, J., Grundy, L., Schober, G., Garcia-Caraballo, S., Zhao, T., Herzig, V., King, G.F., Brierley, S.M., and Lewis, R.J. (2021). A spider-venom peptide with multitarget activity on sodium and calcium channels alleviates chronic visceral pain in a model of irritable bowel syndrome. Pain 162, 569–581.

Cardoso, F.C., Dekan, Z., Rosengren, K.J., Erickson, A., Vetter, I., Deuis, J., Herzig, V., Alewood, P., King, G.F., and Lewis, R.J. (2015). Identification and characterization of ProTx-III [μ-TRTX-Tp1a], a new voltage-gated sodium channel inhibitor from venom of the tarantula Thrixopelma Pruriens. Mol Pharmacol 88, 291–303.

Cardoso, F.C., Dekan, Z., Smith, J.J., Deuis, J.R., Vetter, I., Herzig, V., Alewood, P.F., King, G.F., and Lewis, R.J. (2017). Modulatory features of the novel spider toxin μ-TRTX-Df1a isolated from the venom of the spider Davus fasciatus. Br J Pharmacol 174, 2528–2544.

Cardoso, F.C., and Lewis, R.J. (2018). Sodium channels and pain: from toxins to therapies. Br J Pharmacol 175, 2138–2157.

Cardoso, F.C., and Lewis, R.J. (2019). Structure-Function and Therapeutic Potential of Spider Venom-Derived Cysteine Knot Peptides Targeting Sodium Channels. Front Pharmacol 10, 366.

Cardoso, F.C., Pacifico, L.G., Carvalho, D.C., Victoria, J.M., Neves, A.L., Chavez-Olortegui, C., Gomez, M.V., and Kalapothakis, E. (2003). Molecular cloning and characterization of *Phoneutria nigriventer* toxins active on calcium channels. Toxicon 41, 755–763.

Cardoso, F.C., Pineda, S.S., Herzig, V., Sunagar, K., Shaikh, N.Y., Jin, A., King, G.F., Alewood, P., Lewis, R.J., and Dutertre, S. (2022). The deadly toxin arsenal of the tree-dwelling Australian funnel-web spiders. International Journal for Molecular Bioscience.

Cavalli, J., De Assis, P.M., Cristina Dalazen Goncalves, E., Daniele Bobermin, L., Quincozes-Santos, A., Raposo, N.R.B., Gomez, M.V., and Dutra, R.C. (2022). Systemic, intrathecal, and intracerebroventricular antihyperalgesic effects of the calcium channel blocker CTK 01512-2 toxin in persistent pain models. Mol Neurobiol 59, 4436–4452.

Cordeiro Mdo, N., De Figueiredo, S.G., Valentim Ado, C., Diniz, C.R., Von Eickstedt, V.R., Gilroy, J., and Richardson, M. (1993). Purification and amino acid sequences of six Tx3 type neurotoxins from the venom of the Brazilian ‘armed’ spider *Phoneutria nigriventer* (Keys). Toxicon 31, 35–42.

Da Silva, C.N., Dourado, L.F.N., De Lima, M.E., and Da Silva Cunha-Jr, A. (2020). PnPP-19 peptide as a novel drug candidate for topical glaucoma therapy through nitric oxide release. Transl Vis Sci Technol 9, 33.

De Lima, M.E., Stankiewicz, M., Hamon, A., De Figueiredo, S.G., Cordeiro, M.N., Diniz, C.R., Martin-Eauclaire, M., and Pelhate, M. (2002). The toxin Tx4(6-1) from the spider *Phoneutria nigriventer* slows down Na^+^ current inactivation in insect CNS via binding to receptor site 3. J Insect Physiol 48, 53–61.

Diniz, M.R., Theakston, R.D., Crampton, J.M., Nascimento Cordeiro, M., Pimenta, A.M., De Lima, M.E., and Diniz, C.R. (2006). Functional expression and purification of recombinant Tx1, a sodium channel blocker neurotoxin from the venom of the Brazilian “armed” spider, Phoneutria nigriventer. Protein Expr Purif 50, 18–24.

Diniz, M.R.V., Paiva, A.L.B., Guerra-Duarte, C., Nishiyama, M.Y., Jr., Mudadu, M.A., Oliveira, U., Borges, M.H., Yates, J.R., and Junqueira-De-Azevedo, I.L. (2018). An overview of *Phoneutria nigriventer* spider venom using combined transcriptomic and proteomic approaches. PLoS One 13, e0200628.

Dos Santos, R.G., Van Renterghem, C., Martin-Moutot, N., Mansuelle, P., Cordeiro, M.N., Diniz, C.R., Mori, Y., De Lima, M.E., and Seagar, M. (2002). Phoneutria nigriventer ω-phonetoxin IIA blocks the CaV2 family of calcium channels and interacts with ω-conotoxin-binding sites. J Biol Chem 277, 13856–13862.

Emerich, B.L., Ferreira, R.C., Cordeiro, M.N., Borges, M.H., Pimenta, A.M., Figueiredo, S.G., Duarte, I.D., and De Lima, M.E. (2016). δ-Ctenitoxin-Pn1a, a peptide from *Phoneutria nigriventer* spider venom, shows antinociceptive effect involving opioid and cannabinoid systems, in Rats. Toxins (Basel) 8, 106.

Estrada-Gomez, S., Cardoso, F.C., Vargas-Munoz, L.J., Quintana-Castillo, J.C., Arenas Gomez, C.M., Pineda, S.S., and Saldarriaga-Cordoba, M.M. (2019). Venomic, transcriptomic, and bioactivity analyses of Pamphobeteus verdolaga venom reveal complex disulfide-rich peptides that modulate calcium channels. Toxins (Basel) 11.

Garcia Mendes, M.P., Carvalho Dos Santos, D., Rezende, M.J.S., Assis Ferreira, L.C., Rigo, F.K., Jose De Castro Junior, C., and Gomez, M.V. (2021). Effects of intravenous administration of recombinant Phα1β toxin in a mouse model of fibromyalgia. Toxicon 195, 104–110.

Gewehr, C., Oliveira, S.M., Rossato, M.F., Trevisan, G., Dalmolin, G.D., Rigo, F.K., De Castro Junior, C.J., Cordeiro, M.N., Ferreira, J., and Gomez, M.V. (2013). Mechanisms involved in the nociception triggered by the venom of the armed spider Phoneutria nigriventer. PLoS Negl Trop Dis 7, e2198.

Hu, H., Mawlawi, S.E., Zhao, T., Deuis, J.R., Jami, S., Vetter, I., Lewis, R.J., and Cardoso, F.C. (2021). Engineering of a spider peptide via conserved structure-function traits optimizes sodium channel inhibition *in vitro* and anti-nociception in vivo. Front Mol Biosci 8, 742457.

Isbister, G.K., and Fan, H.W. (2011). Spider bite. Lancet 378, 2039–2047.

Joviano-Santos, J.V., Valadao, P.a.C., Magalhaes-Gomes, M.P.S., Fernandes, L.F., Diniz, D.M., Machado, T.C.G., Soares, K.B., Ladeira, M.S., Massensini, A.R., Gomez, M.V., Miranda, A.S., Tapia, J.C., and Guatimosim, C. (2022). Neuroprotective effect of CTK 01512-2 recombinant toxin at the spinal cord in a model of Huntington’s disease. Exp Physiol.

Jumper, J., Evans, R., Pritzel, A., Green, T., Figurnov, M., Ronneberger, O., Tunyasuvunakool, K., Bates, R., Zidek, A., Potapenko, A., Bridgland, A., Meyer, C., Kohl, S.a.A., Ballard, A.J., Cowie, A., Romera-Paredes, B., Nikolov, S., Jain, R., Adler, J., Back, T., Petersen, S., Reiman, D., Clancy, E., Zielinski, M., Steinegger, M., Pacholska, M., Berghammer, T., Bodenstein, S., Silver, D., Vinyals, O., Senior, A.W., Kavukcuoglu, K., Kohli, P., and Hassabis, D. (2021). Highly accurate protein structure prediction with AlphaFold. Nature 596, 583–589.

King, G.F. (2004). The wonderful world of spiders: preface to the special Toxicon issue on spider venoms. Toxicon 43, 471–475.

King, G.F. (2011). Venoms as a platform for human drugs: translating toxins into therapeutics. Expert Opin Biol Ther 11, 1469–1484.

King, G.F., Gentz, M.C., Escoubas, P., and Nicholson, G.M. (2008). A rational nomenclature for naming peptide toxins from spiders and other venomous animals. Toxicon 52, 264–276.

King, G.F., and Hardy, M.C. (2013). Spider-venom peptides: structure, pharmacology, and potential for control of insect pests. Annu Rev Entomol 58, 475–496.

Kono, N., Nakamura, H., Ohtoshi, R., Moran, D.a.P., Shinohara, A., Yoshida, Y., Fujiwara, M., Mori, M., Tomita, M., and Arakawa, K. (2019). Orb-weaving spider *Araneus ventricosus* genome elucidates the spidroin gene catalogue. Sci Rep 9, 8380.

Kushmerick, C., Kalapothakis, E., Beirao, P.S., Penaforte, C.L., Prado, V.F., Cruz, J.S., Diniz, C.R., Cordeiro, M.N., Gomez, M.V., Romano-Silva, M.A., and Prado, M.A. (1999). Phoneutria nigriventer toxin Tx3-1 blocks A-type K+ currents controlling Ca oscillation frequency in GH3 cells. J Neurochem 72, 1472–1481.

Martin-Moutot, N., Mansuelle, P., Alcaraz, G., Dos Santos, R.G., Cordeiro, M.N., De Lima, M.E., Seagar, M., and Van Renterghem, C. (2006). Phoneutria nigriventer toxin 1: a novel, state-dependent inhibitor of neuronal sodium channels that interacts with μ-conotoxin binding sites. Mol Pharmacol 69, 1931–1937.

Nunes Da Silva, C., Nunes, K.P., De Marco Almeida, F., Silva Costa, F.L., Borges, P.V., Lacativa, P., Pimenta, A.M.C., and Elena De Lima, M. (2019). PnPP-19 peptide restores erectile function in hypertensive and diabetic animals through intravenous and topical administration. J Sex Med 16, 365–374.

Paiva, A.L., Matavel, A., Peigneur, S., Cordeiro, M.N., Tytgat, J., Diniz, M.R., and De Lima, M.E. (2016). Differential effects of the recombinant toxin PnTx4(5-5) from the spider *Phoneutria nigriventer* on mammalian and insect sodium channels. Biochimie 121, 326–335.

Pedron, C., Antunes, F.T.T., Rebelo, I.N., Campos, M.M., Correa, A.P., Klein, C.P., De Oliveira, I.B., Do Nascimento Cordeiro, M., Gomez, M.V., and De Souza, A.H. (2021). Phoneutria nigriventer Tx3-3 peptide toxin reduces fibromyalgia symptoms in mice. Neuropeptides 85, 102094.

Peigneur, S., De Lima, M.E., and Tytgat, J. (2018). Phoneutria nigriventer venom: A pharmacological treasure. Toxicon 151, 96–110.

Pipkorn, R., Boenke, C., Gehrke, M., and Hoffmann, R. (2002). High-throughput peptide synthesis and peptide purification strategy at the low micromol-scale using the 96-well format. J Pept Res 59, 105–114.

Pymol The PyMOL Molecular Graphics System, Version 2.0 Schrödinger, LLC.

Raposo, C., Bjorklund, U., Kalapothakis, E., Biber, B., Alice Da Cruz-Hofling, M., and Hansson, E. (2016). Neuropharmacological effects of *Phoneutria nigriventer* venom on astrocytes. Neurochem Int 96, 13–23.

Richardson, M., Pimenta, A.M.C., Bemquerer, M.P., Santoro, M.M., Beirao, P.S.L., Lima, M.E., Figueiredo, S.G., Bloch, C., Jr., Vasconcelos, E.a.R., Campos, F.a.P., Gomes, P.C., and Cordeiro, M.N. (2006). Comparison of the partial proteomes of the venoms of Brazilian spiders of the genus Phoneutria. Comp Biochem Physiol C Toxicol Pharmacol 142, 173–187.

Smith, J.J., Herzig, V., King, G.F., and Alewood, P.F. (2013). The insecticidal potential of venom peptides. Cell Mol Life Sci 70, 3665–3693.

Smith, J.J., Lau, C.H.Y., Herzig, V., Ikonomopoulou, M.P., Rash, L.D., and King, G.F. (2015). “Therapeutic Applications of Spider-Venom Peptides,” in Venoms to Drugs: Venom as a Source for the Development of Human Therapeutics. The Royal Society of Chemistry), 221–244.

Turchetto, J., Sequeira, A.F., Ramond, L., Peysson, F., Bras, J.L., Saez, N.J., Duhoo, Y., Blemont, M., Guerreiro, C.I., Quinton, L., De Pauw, E., Gilles, N., Darbon, H., Fontes, C.M., and Vincentelli, R. (2017). High-throughput expression of animal venom toxins in *Escherichia coli* to generate a large library of oxidized disulphide-reticulated peptides for drug discovery. Microb Cell Fact 16, 6.

Vieira, L.B., Kushmerick, C., Hildebrand, M.E., Garcia, E., Stea, A., Cordeiro, M.N., Richardson, M., Gomez, M.V., and Snutch, T.P. (2005). Inhibition of high voltage-activated calcium channels by spider toxin PnTx3-6. J Pharmacol Exp Ther 314, 1370–1377.

Von Reumont, B.M., Anderluh, G., Antunes, A., Ayvazyan, N., Beis, D., Caliskan, F., Crnkovic, A., Damm, M., Dutertre, S., Ellgaard, L., Gajski, G., German, H., Halassy, B., Hempel, B.F., Hucho, T., Igci, N., Ikonomopoulou, M.P., Karbat, I., Klapa, M.I., Koludarov, I., Kool, J., Luddecke, T., Ben Mansour, R., Vittoria Modica, M., Moran, Y., Nalbantsoy, A., Ibanez, M.E.P., Panagiotopoulos, A., Reuveny, E., Cespedes, J.S., Sombke, A., Surm, J.M., Undheim, E.a.B., Verdes, A., and Zancolli, G. (2022). Modern venomics-Current insights, novel methods, and future perspectives in biological and applied animal venom research. Gigascience 11.

Wisedchaisri, G., Tonggu, L., Gamal El-Din, T.M., Mccord, E., Zheng, N., and Catterall, W.A. (2021). Structural basis for high-affinity trapping of the Na_V_1.7 channel in its resting state by tarantula toxin. Mol Cell 81, 38–48 e34.

Yonamine, C.M., Troncone, L.R., and Camillo, M.A. (2004). Blockade of neuronal nitric oxide synthase abolishes the toxic effects of Tx2-5, a lethal *Phoneutria nigriventer* spider toxin. Toxicon 44, 169–172.

Zhang, Y., Chen, J., Tang, X., Wang, F., Jiang, L., Xiong, X., Wang, M., Rong, M., Liu, Z., and Liang, S. (2010). Transcriptome analysis of the venom glands of the Chinese wolf spider Lycosa singoriensis. Zoology (Jena) 113, 10–18.

